# Timing and original features of flagellum assembly in trypanosomes during development in the tsetse fly

**DOI:** 10.1101/728964

**Authors:** Moara Lemos, Adeline Mallet, Eloïse Bertiaux, Albane Imbert, Brice Rotureau, Philippe Bastin

## Abstract

*Trypanosoma brucei* exhibits a complex life cycle alternating between tsetse flies and mammalian hosts. When parasites infect the fly, cells differentiate to adapt to life in various tissues, which is accompanied by drastic morphological and biochemical modifications especially in the proventriculus. This key step represents a bottleneck for salivary gland infection. Here we monitored flagellum assembly in trypanosomes during differentiation from the trypomastigote to the epimastigote stage, i.e. when the nucleus migrates to the posterior end of the cell. Three-dimensional electron microscopy (Focused Ion Bean Scanning Electron Microscopy, FIB-SEM) and immunofluorescence assays provided structural and molecular evidence that the new flagellum is assembled while the nucleus migrates towards the posterior region of the body. Two major differences with well known procyclic cells are reported. First, growth of the new flagellum begins when the associated basal body is found in a posterior position relative to the mature one. Second, the new flagellum acquires its own flagellar pocket before rotating on the left side of the anterior-posterior axis. FIB-SEM revealed the presence of a structure connecting the new and mature flagellum and serial sectioning confirmed morphological similarities with the flagella connector of procyclic cells. We discuss potential function of the flagella connector in trypanosomes from the proventriculus. These findings show that *T. brucei* finely modulates its cytoskeletal components to generate highly variable morphologies.

**Author Summary:** *Trypanosoma brucei* is a flagellated parasitic protist that causes human African trypanosomiasis, or sleeping sickness and that is transmitted by the bite of tsetse flies. The complex life cycle of *T. brucei* inside the tsetse digestive tract requires adaptation to specific organs and follow a strictly defined order. It is marked by morphological modifications in cell shape and size, as well organelle positioning. In the proventriculus of tsetse flies, *T. brucei* undergoes a unique asymmetric division leading to two very different daughter cells: one with a short and one with a long flagellum. This organelle is crucial for the trypanosome life cycle as it is involved in motility, adhesion and morphogenesis. Here we investigated flagellum assembly using molecular and 3D Electron Microscopy approaches revealing that flagellum construction in proventricular trypanosomes is concomitant with parasite differentiation. During flagellum growth, the new flagellum is connected to the mature one and rotates around the mature one after its emergence at the cell surface. The sequence of events is different from what is observed in the well-studied procyclic stage in culture revealing different processes governing morphological development. These results highlight the importance to study pathogen development in their natural environment.

## Introduction

*Trypanosoma brucei* is a flagellated parasite responsible for African trypanosomiasis that affects humans and cattle. This parasite is transmitted by the bite of a tsetse fly that itself was infected by ingesting trypanosomes during a blood meal on an infected mammalian host. *T. brucei* exhibits a complex life cycle in which a series of duplication and differentiation processes follow a very ordered progression along the digestive tract of tsetse flies (1–6). When the fly is feeding on an infected mammal, bloodstream trypanosomes are ingested and differentiate into the procyclic form that colonize the posterior midgut (1). Then, procyclic cells migrate from the posterior midgut to the proventriculus (PV, also known as cardia) where they undergo several modifications before reaching the salivary glands. This is a critical step in parasite development and actually represents the major bottleneck for a successful life cycle (7). The only human-infective parasites are called metacyclic trypanosomes and are produced in the salivary glands to be released in the saliva (3–5,8). The successive trypanosome adaptations to these different environments include at least metabolic switching, expression of various stage-specific surface molecules and dramatic changes in morphology (1,3,5,9–11).

The basic cell organization of *T. brucei* is characterized by the presence of a flagellum that is attached to the cell body, a nucleus, a single large and ramified mitochondrion whose genome is concentrated in a structure named kinetoplast (9), which is associated to the basal body of the flagellum through the tripartite attachment complex (12,13). Along their life cycle, trypanosomes exhibit different morphologies classified according to the relative position of the nucleus and the kinetoplast along the antero-posterior axis of the cell. The posterior position of the kinetoplast in relation to the nucleus defines the trypomastigote morphology, whereas the anterior position defines the epimastigote morphology (14).

The flagellum is a cylindrical microtubule-based structure composed of the basal body made of 9 microtubule triplets followed by the transition zone (TZ) (9 microtubule doublets) and the axoneme that displays a classical 9+2 structure with nine microtubule doublets and two central singlets. An extra axonemal structure named the paraflagellar rod (PFR) is connected to doublets 4-7 of the axoneme after flagellum emergence from the flagellar pocket (9). The trypanosome cell shape is defined by a corset of subpellicular microtubules found underneath the plasma membrane with the exception of a single region of the body, the flagellar pocket from where the flagellum emerges (9,15,16). The flagellar pocket is an invagination of the plasma membrane forming a bulb-like structure. When trypanosomes enter the cell cycle, one of the morphological hallmarks is the assembly of a new flagellum (15). In procyclic cells, flagellum construction is characterized by the maturation of an existing pro-basal body followed by the elongation of a new TZ and axoneme. The new flagellum is assembled in an anterior position relative to the old one. It invades the existing flagellar pocket and its basal body rotates around the mature one ensuring the division of the flagellar pocket (17). The tip of the new flagellum is physically connected to the side of the mature flagellum via a cytoskeletal structure termed flagella connector, a transmembrane junction only present during the formation of the new flagellum (18–21). The connection ensures that the new flagellum follows the same helical path as the mature one and the transmission of cell polarity (18). The flagellum is involved in multiple important functions of the parasite during the tsetse infection cycle, such as migration from the midgut to the foregut, and adhesion to the salivary gland epithelium (1,5,8,22,23). Moreover, it plays a crucial role in cell morphogenesis (24,25). Finally, it could act as potential environmental sensor (5,26,27).

The effective success of *T. brucei* transmission to a new vertebrate host relies solely on the metacyclic form, the last developmental stage in the vector that is released together with the insect saliva during the blood meal (5,8). To produce metacyclic trypanosomes, parasites need to colonize the salivary glands. This is presumably ensured by the short epimastigote form that is issued from an asymmetric division taking place in the PV (3,10,28). The PV is an organ that separates the midgut from the foregut and produces the peritrophic matrix, a structure composed of chitin and glycoproteins acting as a physical and biochemical barrier that protects the epithelium from the potentially toxic effects of bloodmeal and pathogens (29,30).

Drastic morphological modifications are taking place in the PV. Parasites first exhibit the trypomastigote morphology and the nucleus changes from an oval to an elongated shape, the body length increases and the distance between the nucleus and the kinetoplast decreases until the nucleus occupies a more posterior position assuming an epimastigote morphology (10). The epimastigote form enters in a single duplication cycle where a mother cell gives rise to two different daughter cells: a short and a long epimastigote (3,10). In this asymmetric division the long epimastigote inherits the long mature flagellum while the short epimastigote possesses the newly assembled, but much shorter flagellum, around 10-fold (4,10). The ultrastructural characterization of the asymmetrically dividing trypanosomes by scanning electron microscopy (SEM) suggests that a short new flagellum emerges from the same flagellar pocket as the mature one. The flagella are linked via a flagella connector-like structure as demonstrated by a single image of transmission electron microscopy (TEM) (10). However, these approaches faced some limitations: SEM shows the cell topography but the kinetoplast and the nucleus are not visible hence preventing morphotype identification. Furthermore, the new flagellum is detected only when it is visible outside of the flagellar pocket. In the case of TEM, single thin sections rarely show the relative position of the nucleus and the kinetoplast, as well as the presence of the mature and new flagellum. To circumvent these issues, we revisited *T. brucei* differentiation in the PV using immunofluorescence assays to monitor flagellum markers, TEM serial sectioning and a 3D electron microscopy approached called Focused Ion Bean Scanning Electron Microscopy (FIB-SEM) based on the use of the slice-and-view method to have a more global vision of flagellum assembly. Here we show that the assembly of the new flagellum is initiated earlier than previously reported in PV trypanosomes committed to the asymmetric division. Formation of a transition zone and elongation of the flagellum can already be detected in trypomastigote cells. In contrast to procyclic trypanosomes, flagellum construction is characterized by an early basal body segregation and the rapid acquisition of an independent flagellar pocket, followed by a late flagellum rotation when compared with procyclic cells. The flagella connector structure that links the new flagellum to the mature one is present from the early stages of flagellum construction.

## Material and Methods

### *Trypanosoma brucei* strain and tsetse infection

The AnTat 1.1E is a pleomorphic clone of *T. brucei* originated from a bushbuck in Uganda in 1966 (31). Procyclic trypanosomes expressing a cytoplasmic reporter composed of the red-shifted luciferase (PpyRE9H) fused to the TdTomato red fluorescent protein and a Ty1 tag (PpyRE9H/TY1/TdTomato) (32) were grown at 27° C in SDM-79 medium (33) supplemented with 10% (v/v) heat-inactivated foetal calf serum (FCS) and 8 mM glycerol. *Glossina morsitans morsitans* teneral males were fed with medium SDMG enriched with 10 mM glutathione containing 5 × 10^6^ trypanosomes/ml. Flies were starved for at least 24 h before dissection performed at 14 or 21 days after infection. Tsetse were scrutinized under a M165FC stereo microscope (Leica) for fluorescent parasites emitted by the TdTomato protein that is visible through the tsetse fly cuticle (32). A total of 33 flies were dissected and the whole alimentary tract removed and placed on a glass slide containing a drop of PBS for immunofluorescence or in cold cacodylate buffer for Electron Microscopy (EM) experiments.

### Immunofluorescence assay

Proventriculi of 12 infected flies were placed in a single 100 µl drop of PBS (pH 7.6) on poly-L-lysine coated slides (J2800AMMZ; Thermo Fisher Scientific). The PVs were dilacerated using two 26 gauge needles to release trypanosomes. Cells were left for 5 min to settle before fixation in cold methanol for 10 sec followed by a rehydration step in PBS for 15 min. For immunodetection, slides were incubated for 1 hour at 37° C in 0.1% BSA in PBS with anti-FTZC 1:500 (rabbit polyclonal), which recognizes a protein termed flagellum transition zone component localized in the transition zone (34) and with mAb25 1:10 (IgG2a) a mouse monoclonal antibody, which recognizes the axonemal protein TbSAXO1 (35,36). After three 5-min washes in PBS, slides were incubated for 1 h at 37° C with anti-mouse antibodies coupled to Cy5 (Jackson) and with anti-rabbit antibodies coupled to Alexafluor-488 (Invitrogen) diluted 1:500 in PBS. Slides were washed three times for 5 min in PBS and DNA was labeled with DAPI (10 µg.mL−1). Slides were mounted in ProLong antifade (Invitrogen) and analyzed with a DMI4000 microscope (Leica), objective 100x 1.4 NA and images were acquired with an ORCA-03G camera (Hamamatsu). Image acquisition was performed using Micromanager software.

### Electron microscopy

For EM sample preparation, the entire digestive tracts of 11 flies were placed in a drop of 0.1 M cacodylate buffer (pH 7.2) and fixed in 2.5% glutaraldehyde (Sigma-Aldrich), 4% paraformaldehyde. Entire proventriculi were then separated from the digestive tract and transferred to 1.5 ml Eppendorf tubes containing 500 µl of 2.5% glutaraldehyde, 4% paraformaldehyde in 0.1 M cacodylate buffer (pH 7.2) for 1 or 2 h at 4° C. Fixed samples were washed three times by the addition of fresh 0.1 M cacodylate buffer (pH 7.2) buffer and post-fixed in 1% osmium (EMS) in 0.1 M cacodylate buffer (pH 7.2) enriched with 1.5% potassium ferrocyanide (Sigma-Aldrich) for 50 min in the dark under agitation. Samples were gradually dehydrated in acetone (Sigma-Aldrich) series from 50% to 100%. Proventriculi were oriented along longitudinal or transversal axes and embedded in PolyBed812 resin (EMS) hard protocol (37), followed by polymerization for 48 h at 60° C. For TEM analysis, 80 nm-thick serial sections were post-stained with 10% uranyl acetate followed by 3% lead citrate, and observed in a FEI Tecnai T12 120kV. For FIB-SEM analysis, the resin embedded PVs were mounted on aluminum stubs, with the pyramidal surface of the resin block pointing upwards. The block surface was coated with a 20 nm-thick layer gold-palladium in a Gatan Ion Beam Coater 681 sputtering device and in additional 2 nm platinum layer using the gas injection system placed inside the Field Emission Scanning Electron Microscope (FESEM) Crossbeam Auriga (Carl Zeiss) workstation microscope chamber. The specimen stage was tilted at 54° with 5 mm working distance of the pole piece, at the coincidence point of the electron and the galion beams. The milling conditions for the trench that allowed the view of the cross-section were 10 nA at accelerating voltage of 30kV. The fine polishing of the surface block was performed with 5 nA at 30kV. For the slice series, 1 nA milling current was applied removing a 10 nm layer from the specimen block surface. Scanning EM images were recorded with an aperture of 60 μm in the high-current mode at 1.5kV of the in-lens EsB detector with the EsB grid set to −300 to −500V. The voxel size was 10 nm in x, y, and z. The contrast of back-scattered electron images was inverted and acquired using ATLAS 5 software (Carl Zeiss).

### Data processing, 3D reconstruction and Flagellum measurement

Alignment of stacks was done with the open source software ImageJ (National Institutes of Health (38) and the Amira software was used for visualization (v6.0.1; FEI; Thermo Fisher Scientific). Segmentation and 3D reconstructions were performed manually using Amira software and a color code was attributed as follows: the new flagellum in orange, the mature one in red, the kinetoplast in purple and the nucleus in blue. The new flagellum of trypanosomes was measured by segmenting the first slice of the basal plate up to the tip of the flagellum including the flagellar membrane.

## Results

### Flagellum construction is initiated in trypomastigote parasites from the proventriculus

Infected tsetse flies were dissected and trypanosomes found in the proventriculus were inspected by using different approaches such as immunofluorescence assays (IFA), classical Transmission Electron Microscopy (TEM) and 3D FIB-SEM. As IFA markers, we used an antibody against the flagellum transition zone component FTZC (34) and the monoclonal antibody mAb25 to detect an axoneme microtubule associated protein called TbSAXO1 (35,36) (Fig 1). In the PV, cell types were defined by their nucleus/kinetoplast DNA content (10) and subcategories were discriminated according to the nucleus morphology in oval (Fig 1A and B) and elongated (Fig 1C and D) as visualized by DAPI staining. Figs 1A shows a trypanosome containing an oval nucleus and a single flagellum with one TZ (magenta) and one axoneme (green). However, 24% (n = 16) of trypomastigotes with an oval nucleus possessed a second spot for FTZC (Fig 1B) which is lateral and in close proximity to the mature one (Fig 1B). A trypomastigote containing an elongated nucleus and a flagellum with one TZ and one axoneme is shown in Fig 1C. A second fluorescent signal for FTZC was also observed in trypomastigotes with elongated nucleus (Fig 1D). Cells containing a second fluorescent signal for FTZC represented 54% of this population (n = 60) (Fig 1D). In 21% of these trypanosomes, a tiny mAb25 signal was observed following the FTZC signal (Fig 1D). The increase in distance between the two signals for FTZC and a more posterior position of the new flagellum in relation to the mature one is presumably due to migration of the basal body of the new flagellum (Fig 1D). The length of the mAb25 signal increased in trypanosomes where the nucleus was migrating towards the posterior end during the differentiation process from trypomastigote to epimastigote (Fig 1E). When the nucleus was positioned at the level of the kinetoplast, cells could no longer be defined as trypomastigote or epimastigote, hence they have been termed “transition forms”. In 100% of transition forms cells a second signal for the TZ and for mAb25 were observed. Cells with positive signal for both FTZC and mAb25 signal in different morphotypes have been quantified (Suppl. Fig 1).

**Fig 1.**
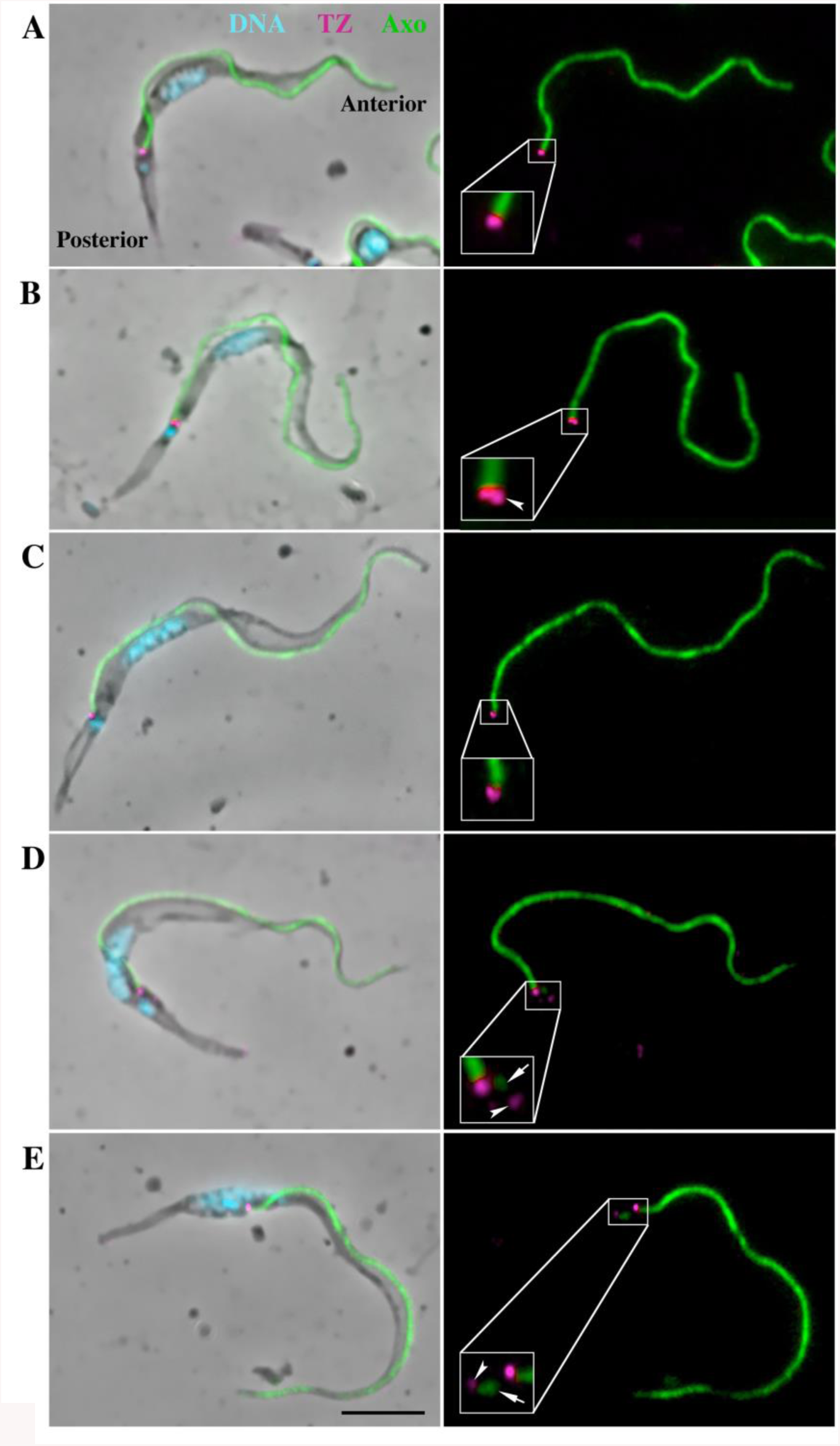
Flagellum assembly is an early event during *T. brucei* differentiation in the PV. (A-E) Immunofluorescence assay using anti-FTZC (magenta) and mAb25 (green) antibodies as markers for TZ and Axoneme, respectively. DAPI was used for DNA staining (blue). (A, B) Trypomastigote cell with an oval nucleus, possessing either a single transition zone (TZ) and axoneme (A) or a second TZ (arrowhead) located laterally to the other one (B). (C, D) Trypomastigote cells with an elongated nucleus possessing either a single TZ and axoneme (C) or with a second axoneme (arrow) being assembled from the second TZ (arrowhead). (E) Late transition form, the kinetoplast is laterally disposed and located at the anterior half of the nucleus whereas the TZ of the new flagellum (arrowhead) is more distant from the TZ of the mature flagellum and the axoneme is being extended. Scale bar: 5µm. Axo, axoneme. Anterior and posterior regions of the cell are indicated on panel A.

These results suggest that a new TZ and a new axoneme are already assembled at the trypomastigote stage in parasites found in the PV. Nevertheless, the signal intensity presumably associated to the new flagellum turned out to be weaker compared with the mature flagellum (Fig. 1D/E). Although IFA demonstrates the presence of FTZC and TbSAXO1 proteins, it does not formally prove that the structures are fully assembled. Considering the differences in intensity of the fluorescent signals, it was crucial to verify the ultrastructural organization by electron microscopy techniques.

### The 3D organization of flagellum assembly in trypomastigotes and transition forms from the proventriculus

FIB-SEM combines an ion beam and an electron for achieving a slice-and-view technique. The ion beam promotes micro abrasions on the sample surface exposing a fresh new layer and the electron beam scans over the block surface generating the image. This process is repeated for several micrometers and the collected images generate 3D stacks with a 10 nm resolution in Z-axis. Two stacks of 15.2 µm and 9.7 µm from the same block were analyzed and representative cells were chosen for segmentation and 3D reconstruction.

Entire PVs were immediately fixed after dissection maintaining the location of trypanosomes in their microenvironment and avoiding any contamination with parasites from the midgut and foregut. The processed PVs were oriented along longitudinal or transversal axes during embedding for better accessibility to the lumen where higher concentrations of trypanosomes were found. The assembly of the new flagellum was investigated in these cells. The earliest stage of flagellum duplication was observed in trypomastigotes with an elongated nucleus (Fig. 2 and video 1). Figure 2A shows the ultrastructural organization of a trypomastigote cell containing a short new flagellum closely associated to the mature one. New and mature flagella were segmented including the flagellar membrane. This parasite was selected for 3D reconstruction (Fig. 2B, C and D). The new flagellum (orange) is closely located to the kinetoplast (purple) (Fig. 2A and B), and is composed of a TZ, a 9+0 microtubule doublet structure in continuity with the basal body (Fig. 2 C, white arrowhead), and a short axoneme of 968 nm in length (from the basal plate of the TZ to the tip), which is entirely enclosed by the flagellar pocket (Fig. 2A and B). The basal bodies are close to each other and separated by only 440 nm. The bottom view of the cell allows the visualization of the entire elongated nucleus (blue) that is 6.8 µm long and occupies a large portion of the cell body (Fig. 2C). The most posterior end of the nucleus is separated from the kinetoplast center by 1.8 µm. Figure 2D shows the top surface view of the cell body and confirms that only the mature flagellum (red) emerges from the flagellar pocket (Fig. 2D).

**Fig 2.**
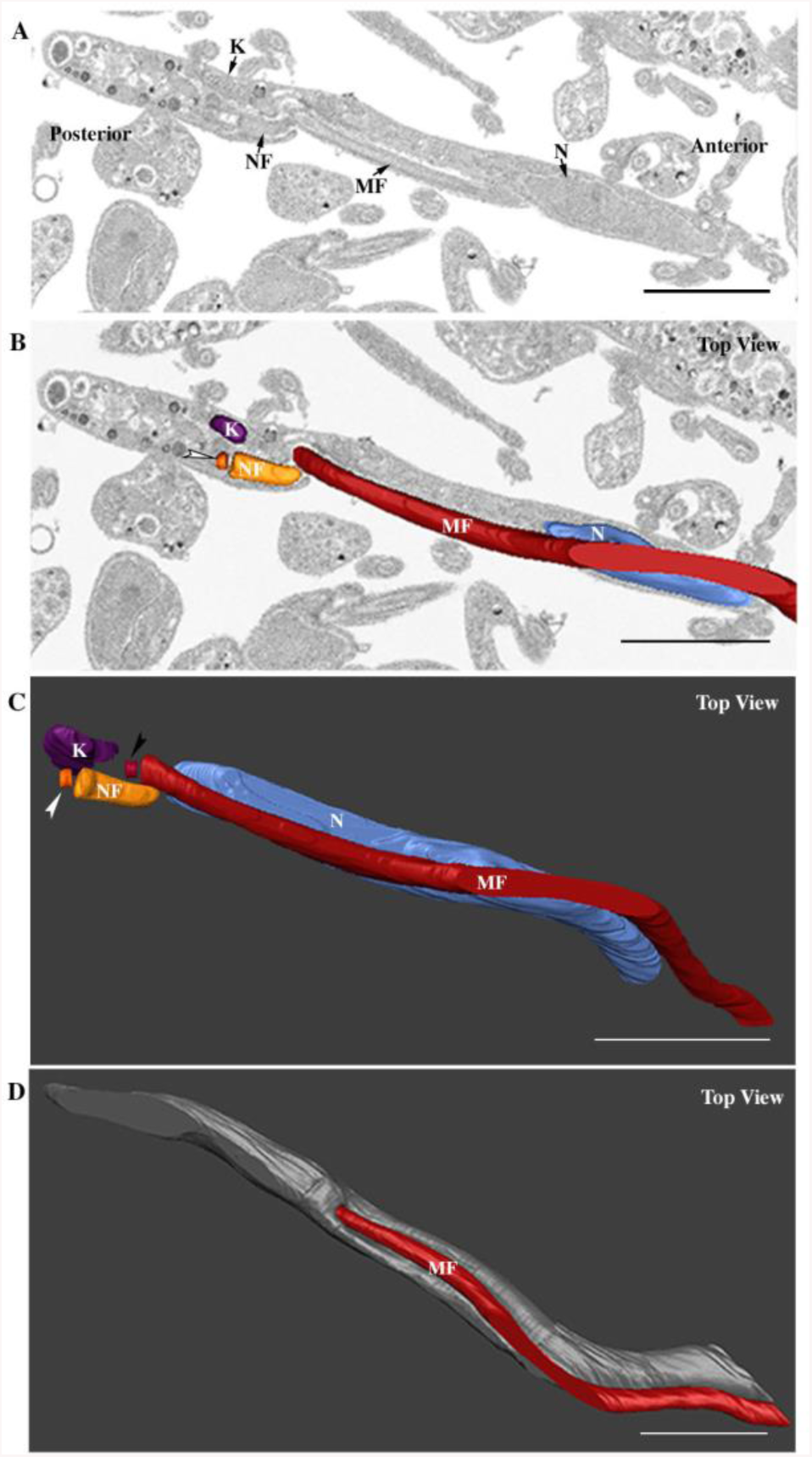
The short new flagellum is inside the flagellar pocket at an early stage of elongation. Images were collected by FIB-SEM and segmented to generate the 3D model. The full stack is shown at video S1. (A) Slice view of a trypomastigote cell, the short new flagellum is laterally connected to the mature flagellum. (B) Top view of segmented kinetoplast (purple), new flagellum (orange) basal body (indicated by a white arrowhead), mature flagellum (red), and nucleus (blue). The new flagellum is located in a more posterior region in relation to the mature one. The diameter of the flagellum looks larger than that of the basal body because the membrane was used for segmentation. (C) Top view of the 3D model showing internal architecture of the cell including the position of the new basal body (white arrowhead) and the mature one (black arrowhead) and the entire elongated nucleus. (D) The new flagellum is inside the flagellar pocket and it is not visualized at the cell surface. The cell body is in grey and the mature flagellum is in red. Scale bars: 2µm. NF, new flagellum; MF, mature flagellum; K, kinetoplast; N, nucleus. White and black arrowheads indicate the new and the mature basal bodies, respectively. Anterior and posterior regions of the cell are indicated on panel A.

Figure 3A shows a section of a trypomastigote cell with the basal body (Fig. 3 C, white arrowhead) of the new flagellum, which is found in a posterior position relative to the mature one (Fig. 3 C, black arrowhead). The basal bodies are separated by 700 nm. The axoneme reaches 1.2 µm and emerges from the flagellar pocket in a posterior position relative to the mature flagellum (Fig. 3B and D). The nucleus is closer to the kinetoplast, as they are separated by only 819 nm (Fig. 3C and video 2).

**Fig 3.**
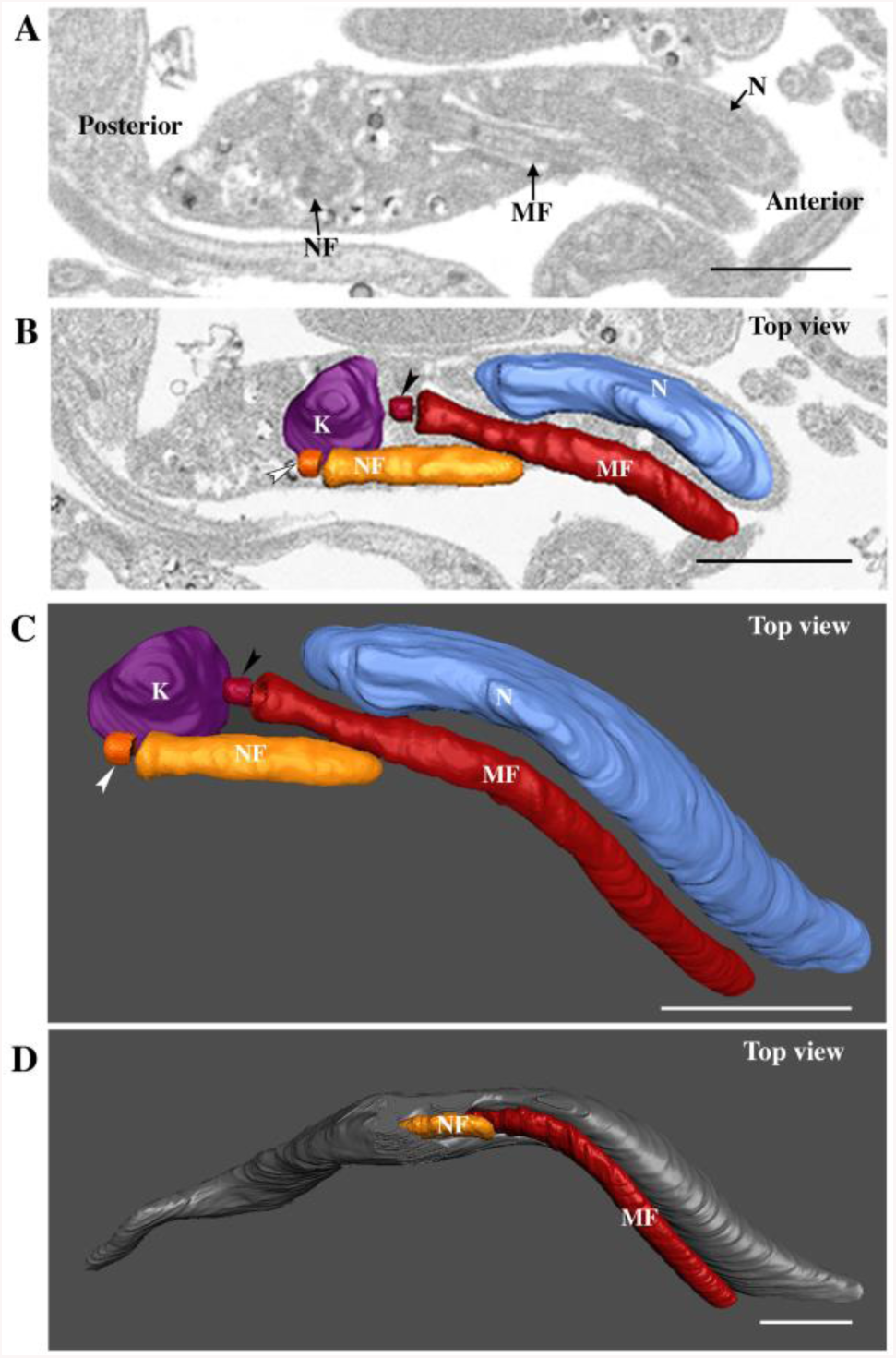
The new flagellum emerges from its flagellar pocket during assembly. The full stack is shown at video S2. (A) Slice view of a trypomastigote cell with the basal body of the new flagellum (arrow), which is posterior in relation to the mature flagellum (MF). (B) Top view of the new flagellum emerging from its own flagellar pocket in a segmented cell. (C) Top view of internal organization of the segmented kinetoplast, new flagellum, mature flagellum and nucleus. The new flagellum is closely associated to the kinetoplast and is in a posterior position in relation with the mature flagellum. (D) Top view of the reconstructed cell body, the new flagellum is now observed at the cell surface. Scale bars: 1µm. NF, new flagellum; MF, mature flagellum; K, kinetoplast; N, nucleus. White and black arrowheads indicate the new and the mature basal bodies, respectively. Anterior region and posterior regions of the cell are indicated on panel A.

For transition forms, parasites were subdivided into early transition form (Fig. 4A-D) and late transition form according to the relative nucleus-kinetoplast position (Fig. 4E-G). In an early transition form, the kinetoplast is enlarged (Fig. 4A) possibly reflecting the duplication of the mitochondrial genome (10). The basal body of the new flagellum is located close to the posterior tip of the kinetoplast (Fig. 4B) and the axoneme measures 2.8 µm. The nucleus and the kinetoplast are occupying the same plane reflecting the moment when the nucleus reaches the posterior tip of the kinetoplast (Fig. 4B and C). This is accompanied by an obvious nucleus deformation (Fig. 4C). Fig. 4D shows the top view of the cell with the new flagellum positioned to the left side relative to the antero-posterior axis of the cell body. A trypanosome in a late transition form is showed in Fig. 4 E-G. The slice view shows the kinetoplast, the nucleus, the new flagellum and the mature flagellum (Fig. 4E). The nucleus migrates towards the posterior region and the kinetoplast is positioned at the anterior half of the nucleus region (Fig. 4 E and F). The basal body of the new flagellum (Fig 4 F, white arrowhead) is positioned at the posterior tip of the kinetoplast, in a similar position as observed in the early transition form (Fig. 4 A-D). However, the new flagellum is laterally positioned to the right side of the antero-posterior axis of the cell body. This is in contrast with all previous stages where the new flagellum is positioned on the left side when looked from the anterior to posterior region of the cell body. The basal bodies are separated by 2.3 µm and the length of the axoneme of the new flagellum reaches 3.7 µm (Fig. 4F and G).

**Fig 4.**
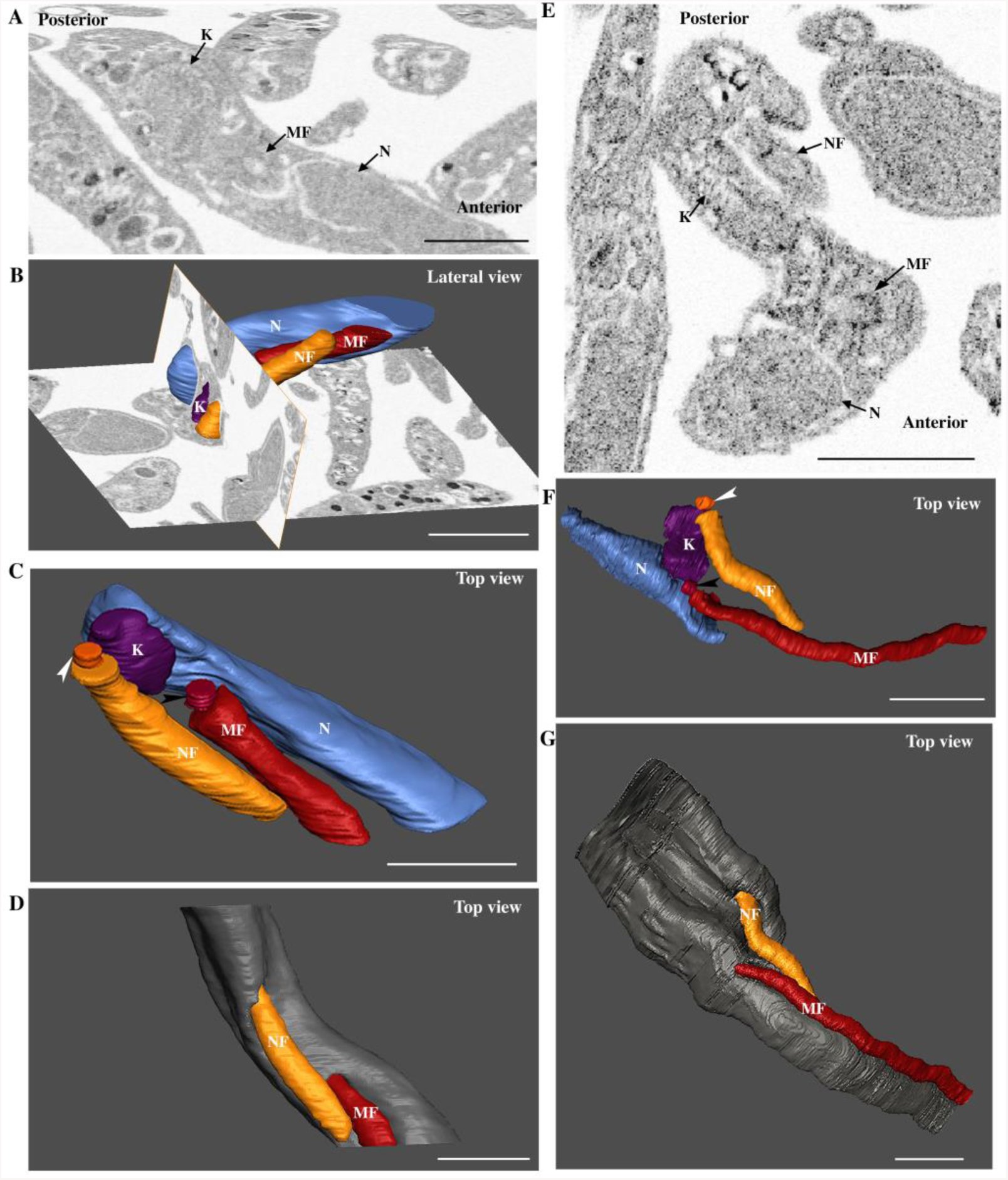
Transition forms where the nucleus migrates towards the posterior region of the body. (A-D) Early transition form and (E-G) late transition form. (A, E) Slice view of trypanosomes exhibiting the nucleus, the kinetoplast, the new flagellum and the mature flagellum. (A) The parasite possesses a large kinetoplast and the new flagellum is not in the plane of the image. (B) Lateral view of the internal organization of a partially reconstructed trypanosome showing that nucleus migration reached the same plane as the kinetoplast. (C, F) 3D model of an early transition form (C) showing nucleus deformation next to the mature flagellum and of a late transition form (F) where the deformation is seen in the region close to the kinetoplast. (D, G) Top view of partially reconstructed trypanosomes. (D) The new flagellum is located to the left side in relation to the mature flagellum. (G) The new flagellum is found at the left side of the posterior-anterior axis of the body. Scale bars: 1µm. NF, new flagellum; MF, mature flagellum; K, kinetoplast; N, nucleus. White and black arrowheads indicate the new and the mature basal bodies, respectively. Anterior region and posterior region of the cell are indicated on panel A and E.

Flagellum elongation in proventricular trypanosomes was measured in cells corresponding to the stages described above (Fig. 5) by taking the first slice of the basal plate up to the tip of the flagellum including the flagellar membrane. The new flagellum elongates progressively, consistent with the order suggested by IFA experiments. When the new flagellum is inside the flagellar pocket, it measures in average 817 nm (n = 10). When this flagellum grows further and can be detected outside of the flagellar pocket in trypomastigote cells, its whole length is 1.5 µm (n = 12), a value that culminates at 2.9 µm (n = 7) in transition forms.

**Fig 5.**
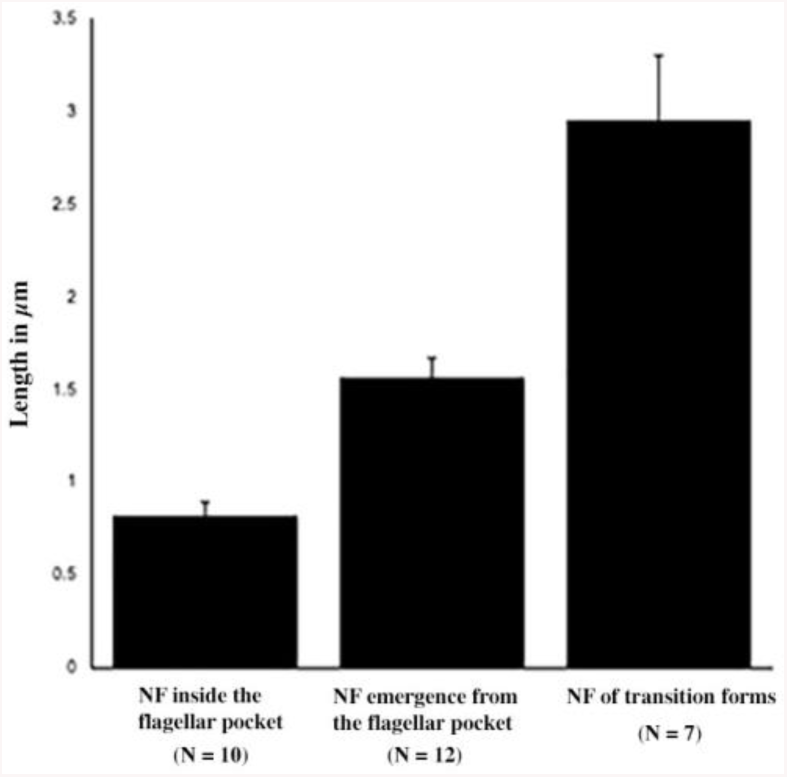
Flagellum elongation in parasites during differentiation in the proventriculus. Measurements of the new flagellum (NF) were taken from the basal plate to the tip of the flagellar membrane in FIB-SEM stacks. Values are given in mean and the SE are indicated. The numbers of parasites is shown in parentheses.

We analyzed 79 trypanosomes during the formation of the new flagellum by using FIB-SEM. The tip of the new flagellum was not visible in 8 cells, but in the 71 other ones an electron-dense structure was observed between the tip of the new flagellum and the side of the mature flagellum. This electron-dense structure seems to connect the new flagellum and the mature one, similar to the flagella connector observed in procyclics (18,19,21). However, the resolution provided by FIB-SEM is not sufficient to analyze its ultrastructure in details. Therefore, we turned to TEM analysis of serial sections, an approach that provides more resolution for structural investigations.

### The new and the mature flagella are associated via a flagella connector structure

Three 80 nm-thick serial sections of a trypomastigote cell with two flagella are shown at Fig. 6. In this cell, a tiny new flagellum is located inside the flagellar pocket, which is shared with the mature flagellum (Fig. 6A-D). This is the earliest stage of flagellum duplication in a trypomastigote cell that could be observed by TEM. The new flagellum consists of a TZ, a basal plate, and a short axoneme (Fig. 6E). A structure linking the new flagellum and the mature one is present and exhibits an electron-dense trilaminate morphology connecting laterally the distal region of the new flagellum with the mature one (Fig. 6C and D – bordeaux arrowheads). Starting from the base of the new axoneme, a first electron-dense plate is facing the basal plate of the new flagellum and the TZ region of the mature one (Fig. 6C – E). The second plate is facing the axoneme of the new flagellum and a portion of the basal plate of the mature one. Finally, the last electron-dense plate is found between the two axonemes.

**Fig 6.**
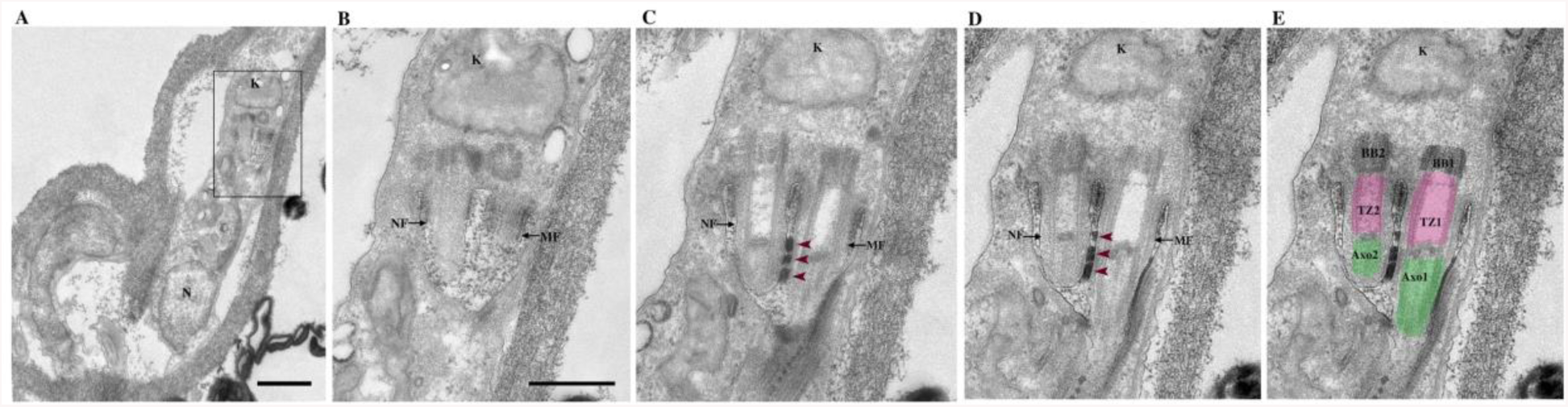
The new flagellum is attached to the mature one via a flagella connector structure. (A) Low magnification of a trypomastigote cell in the PV. (B-D) TEM consecutive 80 nm-thick serial sections of the flagellar pocket region. (E) Image colorized highlighting different structures. (A) A trypomastigote cell with the kinetoplast in a posterior location in relation to the nucleus. (B) The new flagellum shares the flagellar pocket with the mature flagellum. (C, D) The short new flagellum is laterally attached to the mature one via a flagella connector structure (bordeaux arrowheads). The flagella connector is an electron-dense plate structure organized into three distinct layers laterally connecting the distal region of the new flagellum to the mature flagellum. (E) Basal bodies are colorized in dark grey, transition zones in magenta and axonemes in green. BB, TZ and Axo refers to the basal body, transition zone and axoneme, respectively. Number 1 refers to the mature flagellum and number 2 to the new flagellum. Scale bars: 1 µm (A) or 500nm (B-E).

## Discussion

Trypanosome duplication is well documented in procyclic and bloodstream forms. Differentiation from one stage to the other has been well investigated for bloodstream to procyclic conversion (39–41). By contrast, trypanosome development in the tsetse PV is still poorly understood (28). This is explained by the lack of *in vitro* culture for parasites from PV and therefore requires the dissection of a large number of flies which is time-consuming and demands experienced staff.

In this paper, we revisited the process leading to the asymmetric division and the production of long and short epimastigotes in the tsetse proventriculus (Fig. 7). It was thought that the order of the events was first the differentiation of trypomastigotes into epimastigotes that was previously inferred to follow a series of cellular modifications according to a precise chronological plan: (1) nucleus migration to the posterior region of the body, followed by (2) new flagellum assembly, (3) duplication of the kinetoplast and (4) mitosis, ending with (5) cytokinesis to produce one long and one short epimastigote cells (3,10,28). Here, we show that the assembly of the new flagellum is rather initiated before nucleus migration in trypomastigotes (Fig. 7 step 1). The presence of a short new flagellum was unambiguously demonstrated by using specific molecular markers of the TZ (FTZC) and of the axoneme (TbSAXO1) (34–36) combined with 3D FIB-SEM data and TEM serial sections showing the structure of the basal body, the TZ and the axoneme of the new flagellum. First, the short new flagellum is assembled and invades the existing flagellar pocket (Fig. 7 steps 1). This is supported by molecular evidence with a second FTZC signal which is localized in the TZ in trypomastigote parasites with an oval nucleus. Detection of TbSAXO1 indicates that the new flagellum is in a more advanced stage of assembly and correlates with a morphological change in the nucleus from oval to elongated. Such cells types could not have been detected in previous studies because flagellum specific markers were not included (3,10). By DAPI staining and by EM approaches, only one kinetoplast could be observed (10), hence leading to an underestimation of the number of cells that already have initiated assembly of a short new flagellum. Consistent with molecular data, integrated analyses of FIB-SEM results and TEM serial sections provided support for the following series of events (Fig. 7 steps 2 - 5): the new flagellum grows connected to the mature flagellum, the basal bodies segregate and the new flagellum rotates in relation to the mature one. These events take place while the nucleus migrates towards the posterior region of the body (Fig. 7 steps 2 - 5). Although the construction of a new flagellum could be detected by routine EM techniques, evidence for new flagellum assembly can only be obtained if the new flagellum emerges from the flagellar pocket in the case of SEM or when a second flagellum is detected in the same section as the mature one in the case of TEM.

**Fig 7.**
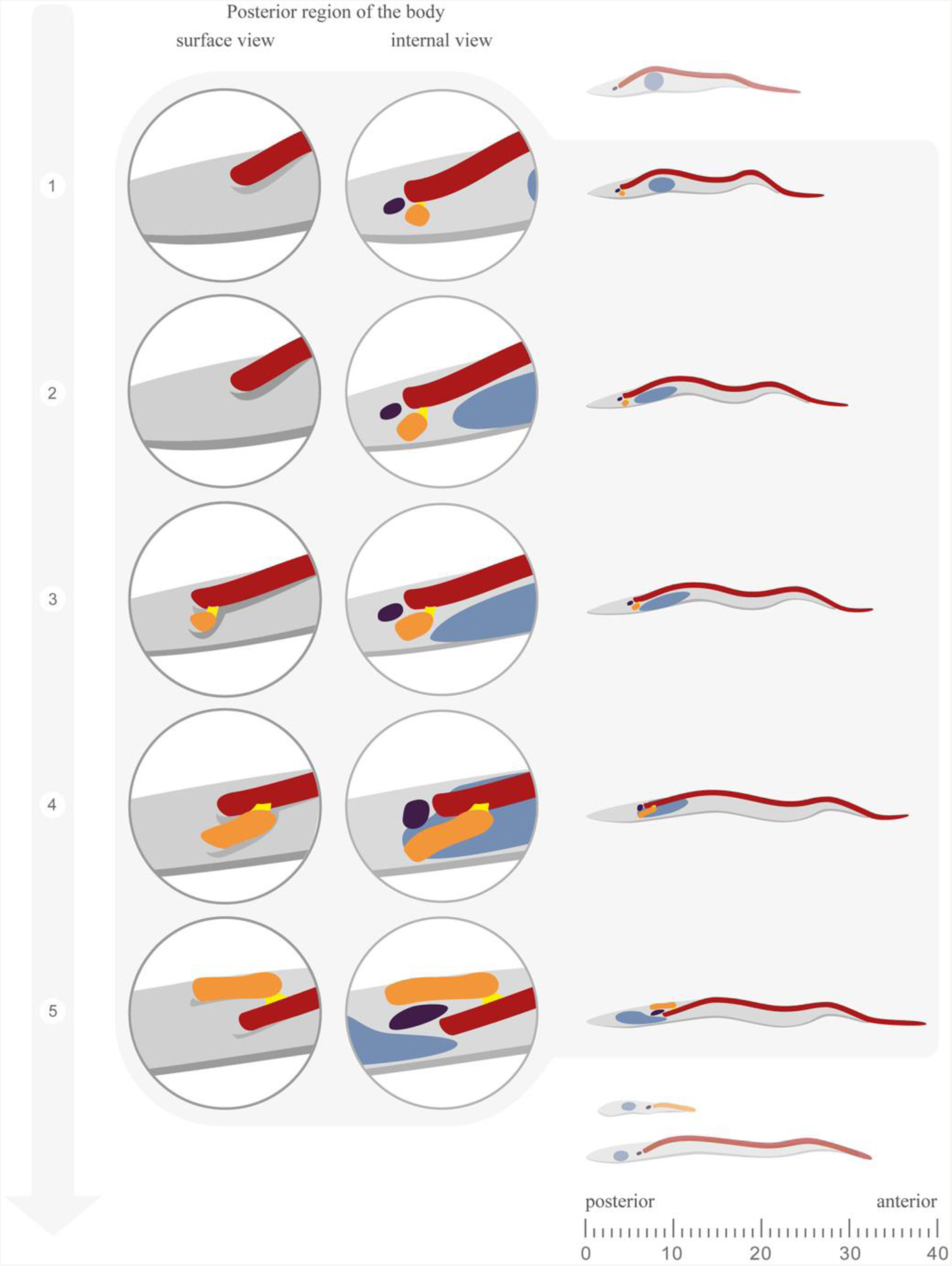
Cartoon illustrating flagellum assembly in trypomastigote cells and in transition forms from the PV. From left to right the first zoom with trypanosomes in dark grey represents the surface view of the posterior region of the body, the emergence of the flagellum from the flagellar pocket whilst the second zoom with trypanosomes in light grey represents the internal view showing the full flagellum elongation process. The new flagellum is represented in orange, the mature flagellum in red, the kinetoplast in purple and the nucleus in blue. At the top, a trypomastigote containing a round nucleus illustrates the stage that precedes the flagellum assembly. On top, the outside view shows a trypanosome where only the mature flagellum is visible outside of the cell body. At stage 1, the internal view represents the 24% of 2F1K1N trypomastigote cells with an oval nucleus and two signals for FTZC in IFA experiments. At stage 2, the short new flagellum is located inside the flagellar pocket whilst only the mature one is visualized outside of the cell body. The internal view represents the 2F1K1N configuration of trypomastigote cells with an elongated nucleus, a new TZ and a new axoneme. The short new flagellum is linked by its tip to the side of the mature one via the flagella connector as demonstrated by serial TEM sectioning. At stage 3, the new flagellum is associated to the mature flagellum via the flagella connector and elongates so that it is observed outside of the cell body. The internal view shows that the nucleus is closer to the kinetoplast indicating the migration towards the posterior region. Stage 4 are early transition form cells where the new flagellum is elongating as demonstrated in the outside view. In the internal view, the nucleus and the kinetoplast are at the same plane. Stage 5 correspond to the late transition form. The outside view shows that the new flagellum has rotated in relation to the mature one. The internal view shows nucleus migration is more advanced in a late transition form. At the bottom, a short and a long epimastigote cell are represented, resulting from the asymmetric division.

The early stages of flagellum construction in PV parasites exhibit differences in the sequence of events when compared to procyclic trypanosomes (17,42). The assembly of the new flagellum in procyclic trypanosomes is initiated in an anterior position relatively to the mature flagellum. The pro-basal body matures into a basal body from which the TZ is assembled. The nascent TZ is adjacent to the mature flagellum and the axoneme elongates while it is still connected to the mature flagellum through the flagella connector (17,19,21,34). The new flagellum undergoes an anticlockwise rotation around the mature one while the basal bodies are almost at the same plane in the antero-posterior axis of the body (17). By contrast, in trypanosomes from the PV, the new flagellum is located in a posterior position in relation to the mature basal body. The distance between the basal bodies increases from 90 nm up to 700 nm and the new flagellum is then observed in the right side of the mature one, suggesting a rotational movement.

The difference in the sequence of events related to basal body segregation and rotation of the flagellum have implications on flagellar pocket morphogenesis. In procyclic trypanosomes, the rotational movement of the basal body facilitates flagellar pocket division (17). In PV trypanosomes, the later rotational movement is observed when the new flagellum has already emerged from its new flagellar pocket indicating that the separation of the flagellar pocket is probably not dependent of the rotational movement of the flagellum.

In all trypanosomes that exhibited two flagella observed by FIB-SEM, an electron-dense structure located at the tip of the new flagellum connecting the lateral aspects of both flagella was detected. TEM serial sections showed that this structure looks similar to the flagella connector described in procyclic trypanosomes (19,21,43). The flagella connector is a three-layered transmembrane junction that joins the tip of the new flagellum to the side of the mature one and is only present during the duplication cell cycle in procyclic trypanosomes. However, it is absent in bloodstream trypomastigotes (18,19,44). A flagella connector structure has previously been observed in trypanosomes from PV by TEM (10). However, the morphotype could not be determined because the nucleus and the kinetoplast were not in the plane of the section.

The flagella connector structure of PV trypanosomes is detected at the early stages of flagellum assembly and persists to the late transition forms. We hypothesize that the flagella connector could be involved in basal body segregation by anchoring the tip of the new flagellum to the mature one in early stages of flagellum elongation when it is inside the flagellar pocket. The flagella connector is facing the region of the basal plate and the proximal region of the axoneme of the mature flagellum and this location is maintained during elongation of the new flagellum until it emerges from the flagellar pocket, which coincides with basal body migration to the posterior region of the body. The stationary position of the flagella connector could prevent further movement towards the anterior region hence elongation of the new flagellum could contribute in moving its basal body to a more posterior position. Such a potential function is consistent with the presence of the basal bodies at opposing poles of the kinetoplast. This is in contrast with procyclic cells where the flagella connector moves towards the mid portion of the old flagellum during elongation until it reaches a position at around 12 µm from the base. At this stage, no further movement of the connector is observed but basal bodies migrate apart, being separated by 6 µm instead of 2 µm (45,46).

These results show the remarkable adaptation of processes driving flagellum assembly and cell morphology in trypanosomes using different tools such as cytoskeletal modifications, flagella connector positioning and flagellar pocket biogenesis to control and produce different morphologies suited for a specific parasite environment.

## Acknowledgements

We thank Aline Crouzols and Christelle Travaillé for help with tsetse fly infection and dissection; Simon Corroyer Dulmont for the Amira training; Sue Vaughan (Oxford Brookes University, UK) and Derrick Robinson (Bordeaux University, France) for critical reading of the manuscript and Derrick Robinson and Frédéric Bringaud for providing the Mab25 and the anti-FTZC antibodies, respectively. We are grateful to the Ultrastructural Bioimaging facility for access to their equipment.

This work is funded by an ANR grant (ANR-18-CE13-0014), by La Fondation pour la Recherche Médicale (Equipe FRM DEQ20150734356) and by a French Government Investissement d’Avenir programme, Laboratoire d’Excellence “Integrative Biology of Emerging Infectious Diseases” (ANR-10-LABX-62-IBEID). We are also grateful for support for FESEM Zeiss Auriga equipment from the French Government Programme Investissements d’Avenir France BioImaging (FBI, N° ANR-10-INSB-04-01) and from a DIM-Malinf grant from the Région Ile-de-France. E.B. was supported by fellowships from French National Ministry for Research and Technology (doctoral school CDV515) and from La Fondation pour la Recherche Médicale (FDT20170436836). The authors declare no competing financial interests.

## Author contributions

M. Lemos infected and dissected the flies, performed the IFA assays and data analyses, prepared sample for FIB-SEM, acquired, segmented and measured the FIB-SEM data, and wrote the paper; A. Mallet acquired the FIB-SEM data, E. Bertiaux infected and dissected the flies; A. Imbert drew the cartoons; B. Rotureau corrected the manuscript; Philippe Bastin coordinated the project and corrected the manuscript. All authors commented on the manuscript.

## Supplementary material

**Suppl Fig. 1.**
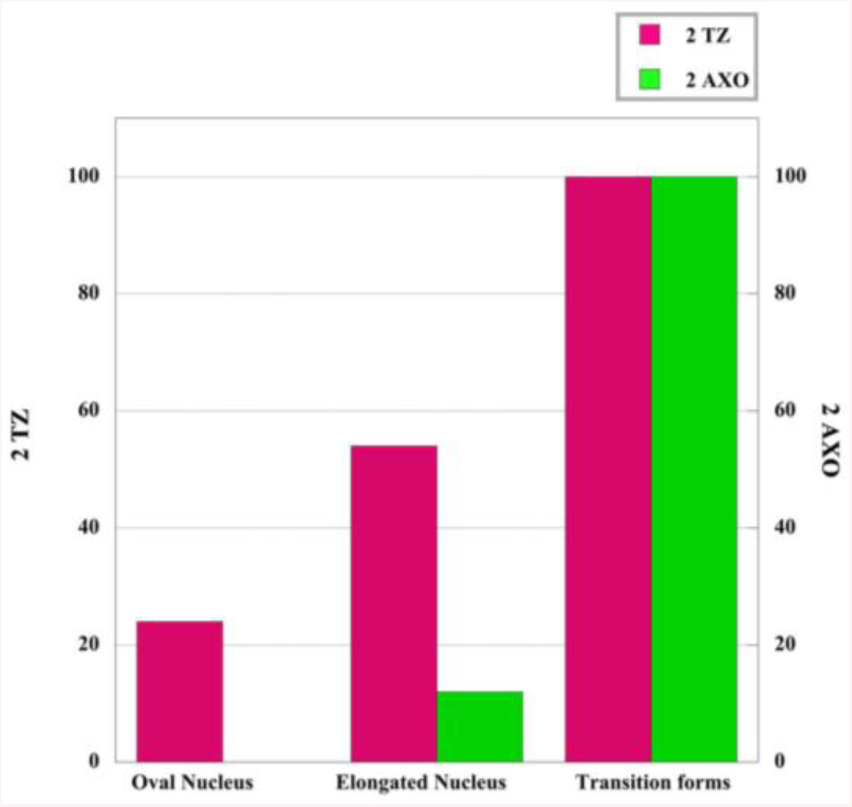
Graph showing the duplication of the TZ and the axoneme in the trypomastigote population from the PV. Scale bar: 5µm. Axo, axoneme. Anterior and posterior regions of the cell are indicated on panel A.

Video 1. 3D reconstruction of a long trypomastigote with a short new flagellum located inside the flagellar pocket.

Video 2. The new flagellum has emerged from its own flagellar pocket and is visible outside of the cell body.

## References

1. Vickerman K. Developmental cycles and biology of pathogenic trypanosomes. Br Med Bull. 1985; 41(2):105–114. https://doi.org/10.1093/oxfordjournals.bmb.a072036.

2. Vickerman K. Developmental cycles and biology of pathogenic trypanosomes. Br Med Bull. 1985; 41(2):105–114. https://doi.org/10.1093/oxfordjournals.bmb.a072036.

3. Van Den Abbeele J, Claes Y, Van Bockstaele D, Le Ray D, Coosemans M. *Trypanosoma brucei* spp. development in the tsetse fly: Characterization of the post-mesocyclic stages in the foregut and proboscis. Parasitology. 1999;118(5): 469–478.

4. Rotureau B, Subota I, Bastin P. Molecular bases of cytoskeleton plasticity during the *Trypanosoma brucei* parasite cycle. Cell Microbiol. 2011;13(5): 705–716. doi: 10.1111/j.1462-5822.2010.01566.x.

5. Rotureau B, Subota I, Buisson J, Bastin P. A new asymmetric division contributes to the continuous production of infective trypanosomes in the tsetse fly. Development. 2012;139(10): 1842–50. doi: 10.1242/dev.072611.

6. Rotureau B, Van Den Abbeele J. Through the dark continent: African trypanosome development in the tsetse fly. Front Cell Infect Microbiol. 2013:53. doi: 10.3389/fcimb.2013.00053.

7. Oberle M, Balmer O, Brun R, Roditi I. Bottlenecks and the maintenance of minor genotypes during the life cycle of *Trypanosoma brucei*. PLoS Pathog. 2010;6(7):1–8. doi: 10.1371/journal.ppat.1001023.

8. Tetley L. Differentiation in *Trypanosoma brucei*: host-parasite cell junctions and theirpersistence during acquisition of the variable antigen coat. J Cell Sci. 1985;74: 1–19.

9. Vickerman K. The mechanism of cyclical development in trypanosomes of the Trypanosoma brucei sub-group: An hypothesis based on ultrastructural observations. Trans R Soc Trop Med Hyg. 1962; 56:487–488. https://doi.org/10.1016/0035-9203(62)90072-X.

10. Sharma R, Peacock L, Gull K, Gluenz E, Gibson W, Carrington M. Asymmetric Cell Division as a Route to Reduction in Cell Length and Change in Cell Morphology in Trypanosomes. Protist. 2008;159(1): 137–151. doi:10.1016/j.protis.2007.07.004.

11. Ooi C-P, Bastin P. More than meets the eye: understanding *Trypanosoma brucei* morphology in the tsetse. Front Cell Infect Microbiol. 2013;3:1–12. doi: 10.3389/fcimb.2013.00071.

12. Robinson DR, Gull K. Basal body movements as a mechanism for mitochondrial genome segregation in the trypanosome cell cycle. Nature 1991; 352 (6337): 731–733. doi: 10.1038/352731a0.

13. Ogbadoyi, E.O; Robinson, D.; Gull K. A High-Order Trans-Membrane Structural Linkage Is Responsible for Mitochondrial Genome Positioning and Segregation by Flagellar Basal Bodies in Trypanosomes. Mol Biol Cell. 2003;13: 3192–3202. doi: 10.1091/mbc.e02-08-0525.

14. Hoare CA, Wallace FG. Developmental Stages of Trypanosomatid Flagellates: a New Terminology. Nature. 1966; 212 (5068): 1385–1386. doi:10.1038/2121385a0.

15. Sherwin T, Gull K. The cell division cycle of *Trypanosoma brucei brucei*: timing of event markers and cytoskeletal modulations. Philos Trans R Soc Lond B Biol Sci. 1989; 323 (1218): 573–588. doi: 10.1098/rstb.1989.0037.

16. Lacomble S, Vaughan S, Gadelha C, Morphew MK, Shaw MK, McIntosh JR, et al. Three-dimensional cellular architecture of the flagellar pocket and associated cytoskeleton in trypanosomes revealed by electron microscope tomography. J Cell Sci. 2009;122 (8): 1081–1090. doi: 10.1242/jcs.045740.

17. Lacomble S, Vaughan S, Gadelha C, Morphew MK, Shaw MK, McIntosh JR, et al. Basal body movements orchestrate membrane organelle division and cell morphogenesis in *Trypanosoma brucei*. J Cell Sci. 2010; 123(17): 2884–2891. doi: 10.1242/jcs.074161.

18. Moreira-Leite FF, Sherwin T, Kohl L, Gull K. A trypanosome structure involved in transmitting cytoplasmic information during cell division. Science. 2001; 294(5542): 610–612. doi: 10.1126/science.1063775.

19. Briggs LJ, McKean PG, Baines A, Moreira-Leite F, Davidge J, Vaughan S, et al. The flagella connector of *Trypanosoma brucei*: an unusual mobile transmembrane junction. J Cell Sci. 2004;117(9): 1641–1651. doi: 10.1242/jcs.00995

20. Vaughan S, Gull K. The structural mechanics of cell division in *Trypanosoma brucei*. Biochem Soc Trans. 2008; 36(3): 421–424. doi: 10.1042/BST0360421.

21. Höög JL, Lacomble S, Bouchet-Marquis C, Briggs L, Park K, Hoenger A, et al. 3D Architecture of the *Trypanosoma brucei* Flagella Connector, a Mobile Transmembrane Junction. PLoS Negl Trop Dis. 2016; 10(1):e0004312. doi: 10.1371/journal.pntd.0004312.

22. Rotureau B, Ooi CP, Huet D, Perrot S, Bastin P. Forward motility is essential for trypanosome infection in the tsetse fly. Cell Microbiol. 2014;16(3):425–33. doi: 10.1111/cmi.12230.

23. Schuster S, Krüger T, Subota I, Thusek S, Rotureau B, Beilhack A, et al. Developmental adaptations of trypanosome motility to the tsetse fly host environments unravel a multifaceted in vivo microswimmer system. Elife. 2017; 6:1–28. doi: 10.7554/eLife.27656.

24. Kohl L, Robinson D, Bastin P. Novel roles for the flagellum in cell morphogenesis and cytokinesis of trypanosomes. EMBO J. 2003; 22(20): 5336–5346. doi: 10.1093/emboj/cdg518.

25. Zhou Q, Liu B, Sun Y, He CY. A coiled-coil- and C2-domain-containing protein is required for FAZ assembly and cell morphology in *Trypanosoma brucei*. J Cell Sci. 2011; 124(22): 3848–3858. doi: 10.1242/jcs.087676.

26. Rotureau B, Morales MA, Bastin P, Späth GF. The flagellum-mitogen-activated protein kinase connection in Trypanosomatids: A key sensory role in parasite signalling and development? Cell Microbiol. 2009;11(5):710–718. doi: 10.1111/j.1462-5822.2009.01295.x.

27. Oberholzer M, Lopez MA, McLelland BT, Hill KL. Social motility in African trypanosomes. PLoS Pathog. 2010; 6(1): e1000739. doi: 10.1371/journal.ppat.1000739.

28. Sharma R, Gluenz E, Peacock L, Gibson W, Gull K, Carrington M. The heart of darkness: growth and form of *Trypanosoma brucei* in the tsetse fly. Trends in Parasito. 2009; (25(11): 517–524. doi: 10.1016/j.pt.2009.08.001.

29. Lehane MJ. Peritrophic matrix structure and function. Annu Rev Entomol. 1997; 42:525–550. doi: 10.1146/annurev.ento.42.1.525.

30. Hegedus D, Erlandson M, Gillott C, Toprak U. New Insights into Peritrophic Matrix Synthesis, Architecture, and Function. Annu Rev Entomol. 2009; 54(1):285–302. doi: 10.1146/annurev.ento.54.110807.090559.

31. Le Ray D, Barry J, Easton C, Vickerman K. first tsetse fly transmission of the AnTat serodeme of *trypanosoma brucei*. Ann Soc Belge Méd Trop. 1977; 57(4–5):369–381.

32. Calvo-Alvarez E, Cren-Travaillé C, Crouzols A, Rotureau B. A new chimeric triple reporter fusion protein as a tool for in vitro and in vivo multimodal imaging to monitor the development of African trypanosomes and Leishmania parasites. Infect Genet Evol. 2018;63:391–403. doi: 10.1016/j.meegid.2018.01.011.

33. Brun R, Schönenberger. Cultivation and in vitro cloning or procyclic culture forms of *Trypanosoma brucei* in a semi-defined medium. Short communication. Acta Trop. 1979; 36(3): 289–292.

34. Bringaud F, Robinson DR, Barradeau S, Biteau N, Baltz D, Baltz T. Characterization and disruption of a new *Trypanosoma brucei* repetitive flagellum protein, using double-stranded RNA inhibition. Mol Biochem Parasitol. 2000; 111(2): 283–297.

35. Pradel LC, Bonhivers M, Landrein N, Robinson DR. NIMA-related kinase TbNRKC is involved in basal body separation in *Trypanosoma brucei*. J Cell Sci. 2006;119(9): 1852–1863. doi: 10.1242/jcs.02900.

36. Dacheux D, Landrein N, Thonnus M, Gilbert G, Sahin A, Wodrich H, et al. A MAP6-Related protein is present in protozoa and is involved in flagellum motility. PLoS One. 2012;7(2): e31344. doi: 10.1371/journal.pone.0031344.

37. Luft JH. Improvements in Epoxy resin embedding methods. J Cell Biol. 1961; 9(2): 409–414. doi: 10.1083/jcb.9.2.409.

38. Schneider CA, Rasband WS, Eliceiri KW. NIH Image to ImageJ: 25 years of image analysis. Nat Methods. 2012; 9(7): 671–675.

39. Matthews KR, Gull K. Cycles within cycles: The interplay between differentiation and cell division in *Trypanosoma brucei*. Parasitol Today. 1994;10(12): 473–476.

40. Tyler KM, Matthews KR, Gull K. Anisomorphic cell division by African trypanosomes. Protist. 2001;152(4): 367–378. doi: 10.1078/1434-4610-00074.

41. Rico E, Rojas F, Mony BM, Szoor B, MacGregor P, Matthews KR. Bloodstream form pre-adaptation to the tsetse fly in *Trypanosoma brucei*. Front Cell Infect Microbio. 2013; 3: 1–15. doi: 10.3389/fcimb.2013.00078.

42. Vaughan S, Gull K. Basal body structure and cell cycle-dependent biogenesis in *Trypanosoma brucei*. Cilia. 2016; 5(1): 1–7. doi: 10.1186/s13630-016-0023-7.

43. Trépout S, Tassin AM, Marco S, Bastin P. STEM tomography analysis of the trypanosome transition zone. J Struct Biol. 2018; 202(1): 51–60. doi: 10.1016/j.jsb.2017.12.005.

44. Hughes L, Towers K, Starborg T, Gull K, Vaughan S. A cell-body groove housing the new flagellum tip suggests an adaptation of cellular morphogenesis for parasitism in the bloodstream form of *Trypanosoma brucei*. J Cell Sci. 2013;126(24): 5748–5757. doi: 10.1242/jcs.139139.

45. Absalon S, Kohl L, Branche C, Blisnick T, Toutirais G, Rusconi F, et al. Basal body positioning is controlled by flagellum formation in *Trypanosoma brucei*. PLoS One. 2007;2(5: e437. doi: 10.1371/journal.pone.0000437.

46. Davidge JA, Chambers E, Dickinson HA, Towers K, Ginger ML, McKean PG, et al. Trypanosome IFT mutants provide insight into the motor location for mobility of the flagella connector and flagellar membrane formation. J Cell Sci. 2006;119(19): 3935–3943. doi: 10.1242/jcs.03203.

